# Sex and metabolic state interact to influence expression of passive avoidance memory in rats: Potential contribution of A2 noradrenergic neurons

**DOI:** 10.1101/2021.02.04.429639

**Authors:** Caitlyn M. Edwards, Tyla Dolezel, Linda Rinaman

## Abstract

Competing motivational drives coordinate behaviors essential for survival. For example, interoceptive feedback from the body during a state of negative energy balance serves to suppress anxiety-like behaviors and promote exploratory behaviors in rats. Results from past research suggest that this shift in motivated behavior is linked to reduced activation of specific neural populations within the caudal nucleus of the solitary tract (cNTS). However, the potential impact of metabolic state and the potential role of cNTS neurons on conditioned avoidance behaviors has not been examined. The present study investigated these questions in male and female rats, using a task in which rats learn to avoid a context (i.e., a darkened chamber) after it is paired with a single mild footshock. When rats later were tested for passive avoidance of the shock-paired chamber, male rats tested in an overnight food-deprived state and female rats (regardless of feeding status) displayed significantly less avoidance compared to male rats that were fed *ad libitum* prior to testing. Based on prior evidence that prolactin-releasing peptide (PrRP)-positive noradrenergic neurons and glucagon-like peptide 1 (GLP1)-positive neurons within the cNTS are particularly sensitive to metabolic state, we examined whether these neural populations are activated in conditioned rats after re-exposure to the shock-paired chamber, and whether neural activation is modulated by metabolic state. Compared to the control condition, chamber re-exposure activated PrRP+ noradrenergic neurons and also activated neurons within the anterior ventrolateral bed nucleus of the stria terminalis (vlBNST), which receives dense input from PrRP+ terminals. In parallel with sex differences in passive avoidance behavior, PrRP+ neurons were less activated in female vs. male rats after chamber exposure. GLP1+ neurons were not activated in either sex. Overnight food deprivation before chamber re-exposure reduced activation of PrRP+ neurons, and also reduced vlBNST activation. Our results support the view that PrRP+ noradrenergic neurons and their inputs to the vlBNST contribute to the expression of passive avoidance memory, and that this contribution is modulated by metabolic state.

## 1. Introduction

Learning to avoid potentially dangerous situations can be essential for survival, and aversive experiences guide future avoidance of those experiences. However, learned avoidance can become excessive and maladaptive, leading to pathological states of anxiety and an increased incidence of stress-related disorders (American Psychiatric Association, 2013; Dymond, 2019; Pittig et al., 2018). Thus, an improved understanding of how the brain controls the expression of learned (i.e., conditioned) avoidance behavior in animal models has potentially high translational value.

It has long been posited that food-seeking/exploratory behaviors compete with avoidance behaviors in a way that is need state-dependent (Hull, 1943). This is evident in early work demonstrating that animals tested in an energy-deficient metabolic state display increased exploratory behavior (Dashiell, 1925), and that food intake is suppressed when animals are exposed to aversive stimuli, a phenomenon called stress-induced hypophagia (Mirsky & Rosvold, 1953). Interestingly, learned/conditioned aversive stimuli also have been demonstrated to suppress food intake (Petrovich, 2009; Petrovich & Lougee, 2011; Reppucci et al., 2013). However, little is known regarding whether metabolic state interacts with avoidance behaviors in response to conditioned aversive stimuli.

The caudal nucleus of the solitary tract (cNTS) is a key circuit node for integrating internal need state with motivated behavior (Maniscalco & Rinaman, 2018). Direct and relayed visceral sensory inputs to the cNTS convey a wide array of interoceptive feedback signals from body to brain, and cNTS neurons relay these signals to the hypothalamus, bed nucleus of the stria terminalis (BNST), and other limbic forebrain regions that control motivated behavior (Bienkowski & Rinaman, 2013; Lin et al., 2002; Maniscalco et al., 2012; Rinaman, 2010). Two distinct neuronal populations in the cNTS, comprising glucagon-like peptide 1 (GLP1)-expressing neurons and the prolactin-releasing peptide (PrRP)-expressing subpopulation of A2 noradrenergic neurons, integrate signals regarding metabolic status with signals of stress generated by real or perceived threats to homeostatic well-being (Holt & Rinaman, submitted; Larsen et al., 1997; Maniscalco & Rinaman, 2018). In rats with *ad libitum* access to food, GLP1+ and PrRP+ neurons are activated to express the immediate-early response gene, cFos, after a variety of innate stressors including restraint (Maniscalco et al., 2015; Maruyama et al., 2001) and exposure to an illuminated elevated platform (Maniscalco et al., 2015). Conversely, when rats are in a state of negative energy balance produced by overnight food deprivation, baseline and stress-induced cFos expression by GLP1+ and PrRP+ neurons is markedly reduced (Edwards et al., 2019; Maniscalco et al., 2015), in conjunction with significant attenuation of “anxiety-like” avoidance behavior (Maniscalco et al., 2015).

The studies cited above support the view that GLP1 and/or PrRP neurons participate in circuits that shape behavioral responses to innate stressors. Less is known regarding the potential role of these neural populations during the expression of conditioned avoidance, in which an animal demonstrates avoidance behavior based on a learned association with an aversive experience, even when the aversive stimulus is no longer present. Two published reports suggest that PrRP+ neurons within the cNTS are involved in the expression of neuroendocrine responses to conditioned fear cues in male rats (Yoshida et al., 2014; Zhu & Onaka, 2003), although GLP1+ neurons were not examined, avoidance behaviors were not examined, and similar studies have not been performed in female rats.

The ventrolateral bed nucleus of the stria terminalis (vlBNST), a downstream target of both PrRP and GLP1 neurons in the cNTS (Bienkowski & Rinaman, 2013; Lin et al., 2002; Maniscalco et al., 2012; Rinaman, 2010), is particularly likely to contribute to passive avoidance behavior. The BNST is broadly implicated in stress and anxiety (Avery et al., 2016; Ch’ng et al., 2018) and has been demonstrated to play a role in contextual fear conditioning (Poulos et al., 2010; Zimmerman & Maren, 2011). Therefore, since acute stress-induced activation of the vlBNST has been shown to be metabolically modulated (Maniscalco et al., 2015), we also investigated conditioned stress-induced neural activation within a specific vlBNST subregion that receives PrRP and GLP1 axonal inputs (i.e., the fusiform nucleus) in rats tested under food-deprived and fed conditions.

The present study tested three hypotheses: (1) metabolic state influences conditioned avoidance behavior, (2) re-exposure to aversive conditioned stimuli activates cNTS GLP1- and PrRP-expressing neurons and neurons in the vlBNST, and (3) food deprivation reduces the ability of conditioned aversive stimuli to activate these neurons. For this purpose, we used a passive avoidance task to examine learned avoidance behavior in rats after a single learning trial. In this task, rats are placed into the illuminated chamber of a light/dark box. The doorway to the darkened chamber is opened, the rat’s latency to enter the darkened chamber is recorded, the doorway is closed, and a single mild electric footshock is then delivered within the darkened chamber. Days later, when the rats are placed again into the light chamber, they remember the aversive experience and avoid entering the dark chamber. The benefit of a single learning trial and a single testing trial permits isolated manipulation of experimental conditions during each phase of the task. To differentiate between the contribution of metabolic state to initial learning and to retrieval and expression of avoidance memory, male and female rats were *ad lib* fed or were food deprived overnight prior to passive avoidance training and/or prior to passive avoidance testing, in a Latin square experimental design. To explore neural correlates of the influence of metabolic state on avoidance behavior, a separate experiment quantified activation of cFos in GLP1+ neurons, noradrenergic neurons and the PrRP+ subset of these neurons, and neurons within the anterior vlBNST in rats re-exposed to the previously shock-paired dark chamber under fed vs. overnight-fasted conditions.

## 2. Methods

All experiments were conducted in accordance with the National Institutes of Health *Guide for the Care and Use of Laboratory Animals* and were reviewed and approved by the Florida State University Animal Care and Use Committee.

### 2.1. Passive Avoidance Task

Adult pair-housed Sprague Dawley rats [Envigo; N=64; 32 males (250-300g body weight), 32 females (150-200g body weight)] were housed in standard tub cages in a temperature- and light-controlled housing environment (lights on from 0400 hr to 1600 hr). Rats were acclimated to handling for three days, with free access to water and rat show (Purina 5001). The day before passive avoidance training (described below), rat pairs were transferred to clean cages with fresh bedding 2 hr before dark onset. Half of the rats (N=16/sex) had continuous free access to chow and water as usual, whereas the remaining rats (N=16/sex) received water but not chow. The following day, after 18-20 hr with no chow access in the food deprived group, all rats underwent passive avoidance training in the light/dark box. Training occurred 4-6 hr after lights on. The light/dark box consisted of a light (illuminated) chamber with clear plastic walls and a smooth white plastic floor, and a dark (non-illuminated) chamber with black plastic walls and a metal grid floor. The two chambers were separated by a metal divider wall with a guillotine door that could be opened and closed remotely. For passive avoidance training, rats were initially individually placed into the light chamber of the box and then the dividing guillotine door was lifted to allow access to the dark chamber. As expected, due to their natural aversion to the light and preference for the dark, rats very quickly entered the dark chamber (average latency = 19.58s; Fig. 1B). Upon entry into the dark chamber, the guillotine door was closed. After a 5s delay, rats received a single mild electric footshock (0.6mA; 1s). Rats remained in the enclosed dark chamber for 30s following footshock and then were returned to their homecage. To reduce the potential influence of food intake immediately after training on experimental outcomes, chow was removed from the cages of *ad lib* fed rats at the time of the learning trial. Chow was returned to all rats (including food-deprived rats) 2 hr prior to dark onset, approximately 6-8 hr after footshock training, to reduce potential effects of post-training food intake on memory consolidation. All rats then remained undisturbed with free access to chow for the next 48 hr.

**Figure 1.**
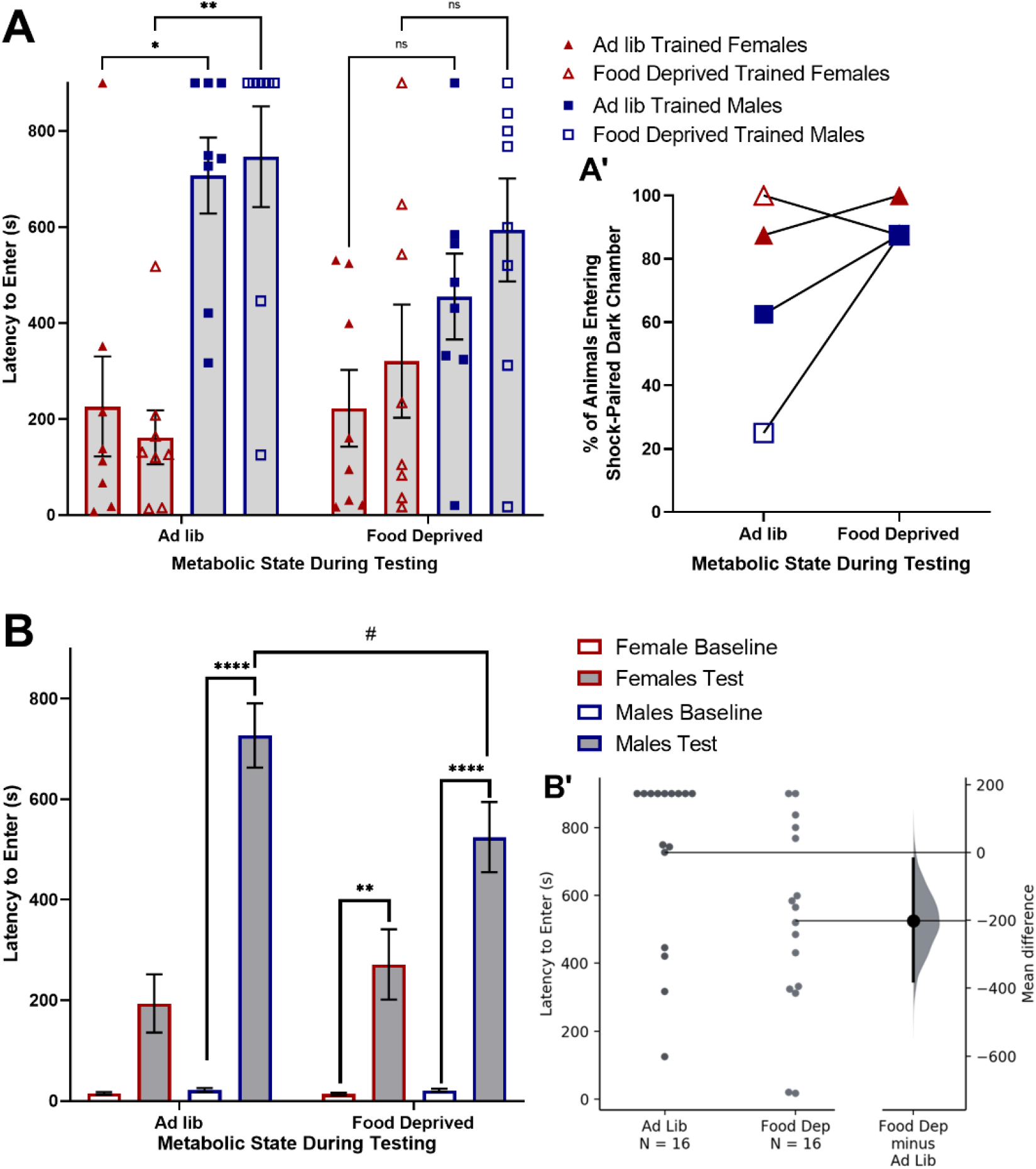
Overnight food deprivation reduces passive avoidance behavior in male but not female rats. (A) Male rats display longer latencies to enter the shock-paired dark chamber compared to female rats when tested under *ad lib* fed conditions, but not when food deprived during testing. There is no effect of food deprivation during training. (A’) The percentage of rats that entered the shock-paired dark chamber during the 900s (15min) test was higher in food deprived males compared to *ad lib* fed males, whereas metabolic state during testing had no effect in females. (B) Data grouped by Metabolic State During Testing (regardless of state during training). Passive avoidance shock training increased latency to enter the dark chamber in both sexes. Food deprivation during testing reduced latency to enter in males, but not females. (B’) Individual data points showing latency to enter the shock-paired dark chamber in male rats that were either *ad lib* fed or food deprived during testing. The mean difference between *ad lib* fed and food deprived male rats with a 95% confidence interval is plotted on a floating axis on the right as a bootstrap sampling distribution. In A and B, ns p > 0.1; #p < 0.1; *p < 0.05; **p < 0.01; ***p <0.001; ****p < 0.0001. Bar graphs in A and B display group mean±SEM.

Two hours before dark onset on the day before testing (i.e., 2 days after training), all rats were transferred to clean cages with chow removed from half of the rats in each of the two original groups (i.e., either *ad lib* fed or food deprived prior to training). This yielded four final experimental groups in a 2×2 design based on metabolic state at training and at testing: (1) Ad lib trained/Ad lib tested; (2) Ad lib trained/Food dep tested; (3) Food dep trained/Ad lib tested; (4) Food dep trained/Food dep tested. On the subsequent testing day (i.e., 3 days after training), rats were tested for passive avoidance memory retention. For this, rats were placed individually into the light chamber of the light/dark box, the guillotine door was opened, and the latency of rats to fully enter the dark chamber was recorded, with a pre-set maximum latency of 900s (15 min). The percentage of rats in each experimental group that entered the dark chamber before 900s elapsed was also determined. During the retention test, the guillotine door remained open and no footshock was administered. After testing, all rats had no access to chow until 2 hr prior to dark onset (i.e., approximately 6-8 hr after testing).

### 2.2. Brain Region Activation

Five to 7 days after the passive avoidance test, a period during which all rats were maintained on *ad lib* chow access, 32 rats (16 males/16 females) were assigned to either a homecage control group (N=16, 8/sex) or were re-exposed to the footshock-associated dark chamber (N=16, 8/sex). Pair-housed rats were each assigned to the same control or re-exposure condition. Rats assigned to the re-exposure group were placed into the dark chamber of the light/dark box for 10 min, 4-6 hr after light onset, with the guillotine door shut; rats were then returned to their homecage. Sixty minutes later, rats were deeply anesthetized with pentobarbital sodium (39 mg/ml i.p., Fatal Plus Solution; Butler Schein) and transcardially perfused with saline (100mL) followed by 4% paraformaldehyde (500mL). Control rats that remained in their homecages were similarly anesthetized and perfused at the same time of day.

A separate group of 24 rats (12 males/12 females) were used to examine the effect of food deprivation on brain activation induced by re-exposure to the footshock-associated dark chamber. Half of these rats (6 males/6 females) were food deprived overnight for 18-20 hr and the other half were fed *ad libitum* before being placed for 10 minutes into the dark chamber of the light/dark box with the guillotine door shut (i.e., all of these rats were re-exposed to the dark chamber). All rats subsequently were returned to their homecages with no access to food, then anesthetized and perfused 60 min later as described above.

### 2.3. Histology

Fixed brains were removed from the skull, post-fixed overnight at 4°C, then cryoprotected in 20% sucrose. Brains were blocked and sectioned coronally (35 μm) using a Leica freezing-stage sliding microscope. Tissue sections were collected in six serial sets and stored at −20°C in cryopreservant solution (Watson et al., 1986) until immunohistochemical processing. Primary and secondary antisera were diluted in 0.1M phosphate buffer containing 0.3% Triton X-100 and 1% normal donkey serum. Two sets of tissue sections from each rat (with each set containing a complete rostrocaudal series of sections spaced by 210 μm) were incubated in a rabbit polyclonal antiserum against cFos (1:10,000; Cell Signaling, 2250; AB_2247211), followed by biotinylated donkey anti-rabbit IgG (1:500; Jackson ImmunoResearch). Sections were then treated with Elite Vectastain ABC reagents (Vector) and reacted with diaminobenzidine (DAB) intensified with nickel sulfate to produce a blue-black nuclear cFos reaction product.

To visualize cFos within hindbrain GLP1 neurons, one set of cFos-labeled tissue sections was subsequently incubated in a rabbit polyclonal antiserum against GLP1 (1:10,000; Bachem, T-4363; AB_518978), followed by biotinylated donkey anti-rabbit IgG (1:500; Jackson ImmunoResearch), Elite Vectastain ABC reagents (Vector), and reacted with plain DAB to produce a brown cytoplasmic reaction product. In another set of cFos-reacted tissue from each rat, a similar dual immunoperoxidase labeling process was used to visualize cFos within a caudal subset of A2 neurons that express prolactin-releasing peptide (PrRP) [rabbit anti-PrRP; 1:1000; Phoenix Pharmaceuticals, H-008-52].

To visualize cFos within all noradrenergic neurons, a third set of tissue was incubated in a cocktail of rabbit polyclonal antiserum against cFos (1:1000, Cell Signaling, 2250; AB_2247211) and a mouse monoclonal antiserum against dopamine beta-hydroxylase (DbH; 1:5000; Millipore, MAB308; AB_2245740), followed by a cocktail of Cy3-conjugated donkey anti-rabbit Cy3 (red) and Alexa Fluor 488-conjugated donkey anti-mouse (green).

### 2.4. Quantification of Neural Activation

The cNTS (~15.46-13.15 mm caudal to bregma) was imaged at 20x using a Keyence microscope (BZ-X700). Using separate sets of dual-labeled tissue sections from each rat, the total numbers of GLP1+, DbH+, and PrRP+ cNTS neurons were counted, and the percentage of each chemically identified neural population co-expressing cFos was determined. A neuron was considered to be cFos-positive if its nucleus contained any visible cFos immunolabeling.

The anterior vlBNST (~0.26-0.40mm caudal to bregma) was imaged at 10x. Using ImageJ, a circular region of interest (ROI) of 100μm^2^ was centered on the area containing the highest density of DbH+ fibers and terminals; this region corresponds to the fusiform subnucleus of the anterior vlBNST, which also contains the highest density of PrRP+ fibers and terminals within the rat brain (Lin et al., 2002; Maniscalco et al., 2015; Maruyama et al., 1999). The number of cFos-positive neurons within each 100μm^2^ ROI was determined bilaterally in 2-3 sections per brain and averaged across assessed ROIs in each animal.

### 2.5. Statistics

Data were analyzed using GraphPad Prism. Passive avoidance behavior was analyzed in three ways: (1) Latency to enter the dark chamber was compared between groups using a 2×2×2 ANOVA with sex, metabolic state during training, and metabolic state during testing as independent variables; (2) Within each sex, the proportion of rats entering the dark chamber before the 900s maximum was compared between groups that were either food deprived or *ad libitum* fed at the time of testing, using a simple logistic regression; (3) Latency to enter the dark chamber was compared between groups using a 2×2×2 ANOVA with shock training, sex, and metabolic state during testing as independent variables. Cell count data were analyzed using a 2×2 ANOVA with either sex and dark chamber exposure or sex and metabolic state as independent variables. When ANOVA F values indicated significant main effects and/or interactions, post-hoc comparisons were made using Tukey’s multiple comparisons tests. An alpha level of 0.05 (p ≤ 0.05) was used as the criterion for statistically significant effects.

## 3. Results

### 3.1. Passive Avoidance Behavior

Between subjects three-way ANOVA (Sex x Metabolic State During Training x Metabolic State During Testing) revealed a significant main effect of sex [F(1,56) = 34.79; p < 0.001] and a significant interaction between Sex and Metabolic State During Testing [F(1,56) = 4.403; p = 0.04] on latency to enter the shock-paired dark chamber (Fig. 1A). Post-hoc Tukey’s multiple comparisons tests indicated that male rats that were *ad libitum* fed during testing took significantly longer to enter the shock-paired dark chamber compared to *ad libitum* fed female rats; this was true both in the group that was also *ad libitum* fed during training (Ad lib trained/Ad lib Tested; p = 0.0142) and in the group that was food deprived before training (Food dep Trained/Ad lib tested; p = 0.0013). Thus, sex and metabolic status at the time of testing each significantly impacted passive avoidance, whereas metabolic status at the time of training did not. There were no significant sex differences in latency to enter the shock-paired dark chamber in either of the two groups that were food deprived before testing (Ad lib trained/Food dep tested and Food dep trained/Food dep tested). Given the preset maximum latency period, some rats never entered the dark chamber during the 900s (15 min) test. Almost all female rats entered the dark chamber within 900s, regardless of metabolic state (Ad lib trained/Ad lib tested = 87.5%; Ad lib trained/Food dep tested = 100%; Food dep trained/Ad lib tested = 100%; Food dep trained/Food dep tested = 100%; Fig. 1A’). Comparatively fewer male rats tested under the fed condition entered the dark chamber (Ad lib trained/Ad lib tested = 62.5%; Food dep trained/Ad lib tested = 25.0%), whereas a greater proportion of male rats entered the dark chamber when tested under the food deprived condition (Ad lib trained/Food dep tested = 87.5%; Food dep trained/Food dep tested = 87.5%). A simple logistic regression was run to examine group differences in the likelihood of Ad lib tested vs. Food dep tested rats to enter the dark chamber before the 900s maximum. The results revealed that male rats tested in the food deprived state were 9 times more likely to enter the shock-paired dark chamber compared to male rats tested under fed conditions [β = 2.197; z = 2.418; p = 0.0156]. Conversely, metabolic state at the time of testing had no impact on the likelihood that female rats entered the dark chamber before the 900s maximum [β = 3.291; z < 0.0001; p > 0.9999].

As expected, within subjects three-way ANOVA (Pre/Post Shock Training x Sex x Metabolic State During Testing) indicated a significant main effect of shock training [F(1,60) = 160.7; p < 0.001], such that rats displayed significantly less avoidance of the dark chamber (i.e., shorter latencies to enter) before vs. after footshock training (Fig. 1B). There also was a significant main effect of sex [F(1,60) = 36.54; p < 0.001], a significant interaction between Sex and Metabolic State During Testing [F(1,60) = 4.423; p = 0.040], and a significant interaction between Shock Training, Sex, and Metabolic State During Testing [F(1,60) = 35.34; p < 0.001]. Post-hoc Tukey’s multiple comparisons tests indicated significant differences between pre- and post-training entry latencies in *ad lib*-tested males (p < 0.001), food deprived-tested males (p < 0.001), and food deprived-tested females (p = 0.005). The entry latencies of male vs. female rats during testing were significantly different under both *ad lib* (p < 0.001) and food deprived (p = 0.004) testing conditions. In addition, males tested while food deprived displayed shorter latencies to enter the shock-paired dark chamber (i.e., less inhibitory avoidance) compared to males tested in the *ad lib* fed state (p = 0.050). Examining the unpaired mean difference between males tested while food deprived vs. *ad lib* fed using estimation statistics (Ho et al., 2019) revealed that fed rats took an average of 194s (3.23 min) longer to enter the shock-paired dark chamber compared to food deprived rats [95%CI 12.7s, 373s] (Fig. 1B’).

Overall, these behavioral results indicate that male rats tested for passive avoidance under *ad libitum* fed conditions display more avoidance of the shock-paired dark chamber than female rats tested under the same condition. Further, food deprivation prior to testing reduced passive avoidance in males to a level comparable to that displayed by females. Conversely, food deprivation prior to testing did not affect passive avoidance in females, and food deprivation prior to training did not impact passive avoidance during testing in either sex.

### 3.2. Impact of Dark Chamber Exposure on Neural Activation in Ad Lib-Fed Rats

#### 3.2.1. cNTS

##### 3.2.1.1. A2 Noradrenergic Neurons

###### *DbH*+ *A2 neurons*

Two-way ANOVA (Sex x Dark Chamber Exposure) revealed a significant main effect of exposure to the shock-paired dark chamber on DbH+ neuron activation across three rostrocaudal levels of the cNTS [F(1,27) = 44.52; p < 0.0001; Fig. 2A] as well as in each specific cNTS level relative to the AP (caudal to the AP: [F(1,27) = 34.95; p < 0.001]; AP level: [F(1,27) = 52.95; p < 0.001]; rostral to the AP: [F(1,12) = 49.24; p < 0.001]; Fig. 2C). Post-hoc Tukey’s multiple comparisons tests confirmed a significant increase in A2 neural activation in rats exposed to the dark chamber vs. homecage controls; this was true in both females (p = 0.0044) and males (p < 0.0001), with no main effect of sex on the overall activation of DbH+ neurons. There was no interaction between dark chamber exposure and sex on DbH+ neural activation across all rostrocaudal levels; however, there was a significant interaction between dark chamber exposure and sex at the cNTS level caudal to AP [F(1,27) = 5.408; p = 0.0278] and at the level of AP [F(1,27) = 7.432; p = 0.0111] but not rostral to AP. Examining the unpaired mean difference between neural activation in dark chamber re-exposed rats vs. homecage controls using estimation statistics (Ho et al., 2019) indicated that 24.8% more DbH+ neurons were activated in male rats re-exposed to the shock-paired dark chamber compared to activation in male homecage controls [95.0%CI 18.9%, 29.6%], and that 15.9% more DbH+ neurons were activated by dark chamber re-exposure in female rats compared to activation in female homecage controls [95.0%CI 7.21%, 26.3%].

**Figure 2.**
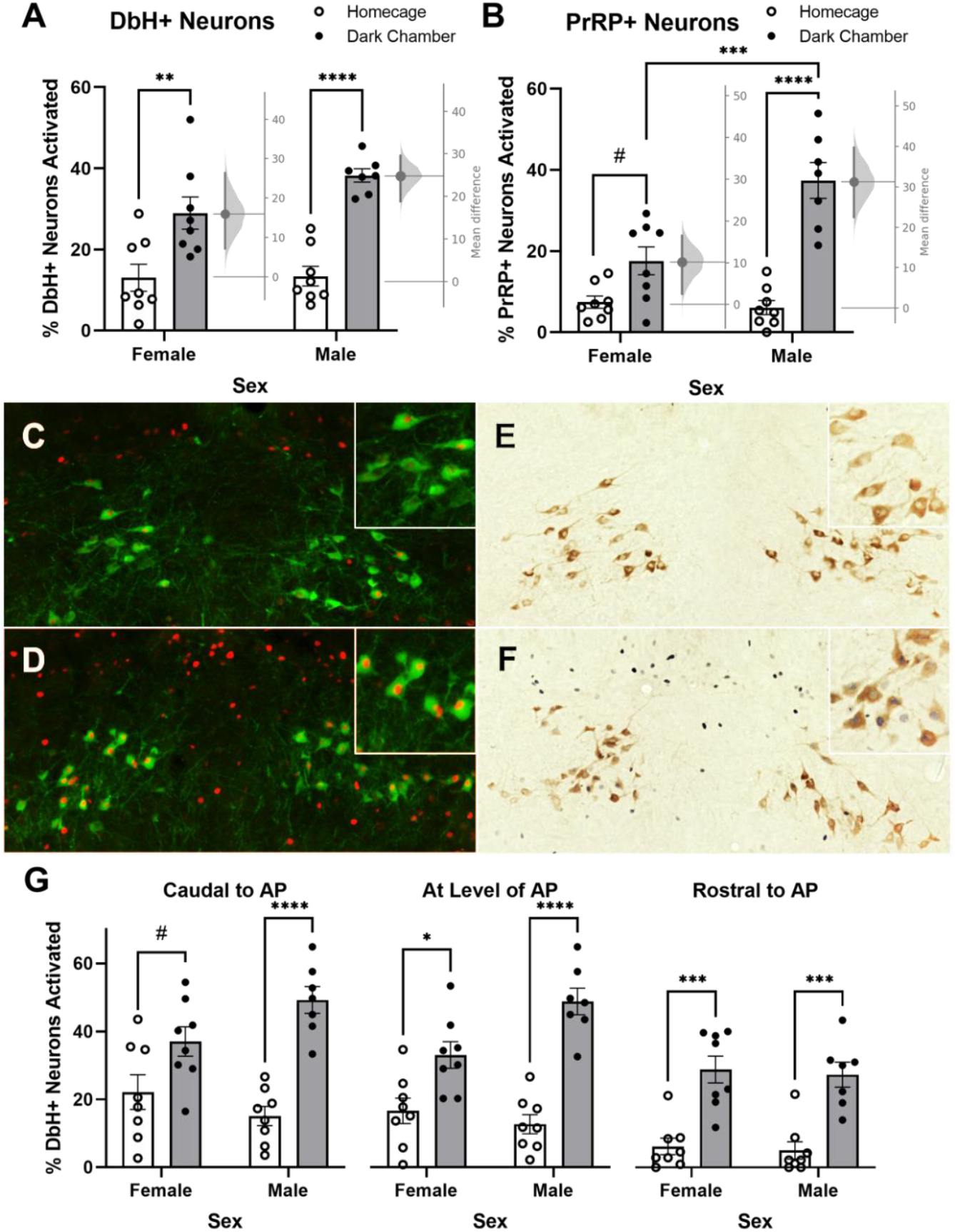
Shock-paired dark chamber re-exposure increases the percentage of noradrenergic neurons in the cNTS expressing cFos. (A) Shock-paired dark chamber re-exposure increases activation of DbH+ neurons in both male and female rats. The mean difference in neural activation between dark chamber re-exposed vs. homecage control female and male rats with a 95% confidence interval is plotted on a floating axis on the right of each set of bars, corresponding a bootstrap sampling distribution for each sex. (B) Shock-paired dark chamber re-exposure increases activation of PrRP+ noradrenergic neurons in male rats, with a trend towards increased activation in female rats. The mean difference in neural activation between dark chamber re-exposed and homecage control female and male rats with a 95% confidence interval is plotted on a floating axis on the right of each set of bars. (C+D) Representative images of nuclear cFos (red) and cytoplasmic DbH (green) immunolabeling in a male homecage control rat (C) and a male dark chamber re-exposed rat (D). Insets in the upper right corner of each panel provide higher-magnification views, showing increased neural activation in the chamber re-exposed rat. (E+F) Representative images of nuclear cFos (dark blue/black) and cytoplasmic PrRP immunolabeling (brown) in a homecage control rat (E) and in a dark chamber re-exposed rat (F). Insets in the upper right corner of each panel provide higher-magnification views, showing increased neural activation in the chamber re-exposed rat. (G) Shock-paired dark chamber re-exposure activates DbH+ noradrenergic neurons in both sexes, although in females the most caudal cNTS level *(left)* displayed only a trend towards increased activation. In A,B, and G, #p < 0.1; *p < 0.05; **p < 0.01; ***p <0.001; ****p < 0.0001. Bars represent group mean±SEM.

###### *PrRP*+ *subpopulation of A2 neurons*

Two-way ANOVA (Sex x Dark Chamber Exposure) revealed a significant main effect of dark chamber exposure [F(1,27) = 50.35; p < 0.0001] and a significant main effect of sex [F(1,27) = 10.00; p = 0.0038] on PrRP+ neuron activation. Additionally, a significant interaction between dark chamber exposure and sex was detected [F(1,27) =13.11; p = 0.0012). Post-hoc Tukey’s multiple comparisons tests confirmed a significant increase in PrRP neural activation between dark chamber re-exposed vs. homecage control male rats (p < 0.0001), with only a trending group difference in females (p = 0.0825). There also was a significant sex difference in PrRP neural activation between dark chamber re-exposed males vs. females (p = 0.0004). Examining the unpaired mean difference in PrRP neural activation between dark chamber re-exposure vs. homecage control conditions using estimation statistics (Ho et al., 2019) indicated that 31.3% more PrRP+ neurons were activated in male rats re-exposed to the dark chamber compared to activation in male homecage controls [95.0%CI 22.5%, 39.7%], and that 10.1% more PrRP+ neurons were activated in female rats re-exposed to the dark chamber compared to activation in female homecage controls [95.0%CI 2.55%, 16.5%].

##### 3.2.1.2. *GLP1*+ *Neurons*

Two-way ANOVA (Sex x Dark Chamber Exposure) indicated a trending main effect of sex [F(1,27) = 3.830; p = 0.0608] but no effect of dark chamber exposure and no interaction between sex and dark chamber exposure on activation of GLP1+ neurons.

#### 3.2.2. BNST

Two-way ANOVA (Sex x Dark Chamber Exposure) revealed a significant main effect of dark chamber exposure on the average number of cFos+ neurons within a 100μm^2^ region of interest (ROI) corresponding to the fusiform subnucleus of the vlBNST, where DbH+ (and PrRP+) axon terminals are most densely clustered [F(1,27) = 29.56; p < 0.0001]. There was no main effect of sex, and no interaction between sex and dark chamber exposure on cFos counts within this region. Post-hoc Tukey’s multiple comparisons tests confirmed that compared to activation under control conditions, dark chamber re-exposure increased neural activation within the anterior vlBNST in both males (p = 0.0024) and females (p = 0.005).

### 3.3. Impact of Metabolic State on Neural Activation After Dark Chamber Exposure

#### 3.3.1. cNTS A2 Noradrenergic Neurons

##### *DbH*+

Two-way ANOVA (Sex x Metabolic State) revealed a significant main effect of metabolic state on DbH+ neuron activation in response to dark chamber re-exposure across all three rostrocaudal levels of the cNTS [F(1,20) = 51.16; p < 0.0001; Fig. 3A], and also specifically at levels caudal to the AP [F(1,20) = 51.21; p < 0.0001] and at the level of the AP [F(1,20) = 19.65; p = 0.0003], but with only a trend towards significance at levels rostral to AP [F(1,20) = 3.430; p = 0.0788; Fig. 3C]. There was also a trend towards a main effect of sex on DbH+ neuron activation across all levels of the cNTS [F(1,20) = 3.684; p = 0.0693], which seems mostly driven by a main effect of sex at the level of the AP [F(1,20) = 5.252; p = 0.0329] and rostral to the AP [F(1,20) = 5.209; p = 0.0336]. There was no interaction between metabolic state and sex on DbH neural activation. Post-hoc Tukey’s multiple comparisons tests indicated that food deprivation prior to dark chamber re-exposure reduced DbH+ neuron activation across all cNTS levels in both males (p < 0.0001) and females (p = 0.0033), and specifically at levels caudal to the AP in both sexes (males p = 0.0001; females p = 0.0009). At the level of the AP, food deprivation reduced DbH+ neuron activation in males (p = 0.0015) but not in females, and there was no effect of metabolic state on DbH+ neural activation rostral to AP in either sex.

**Figure 3.**
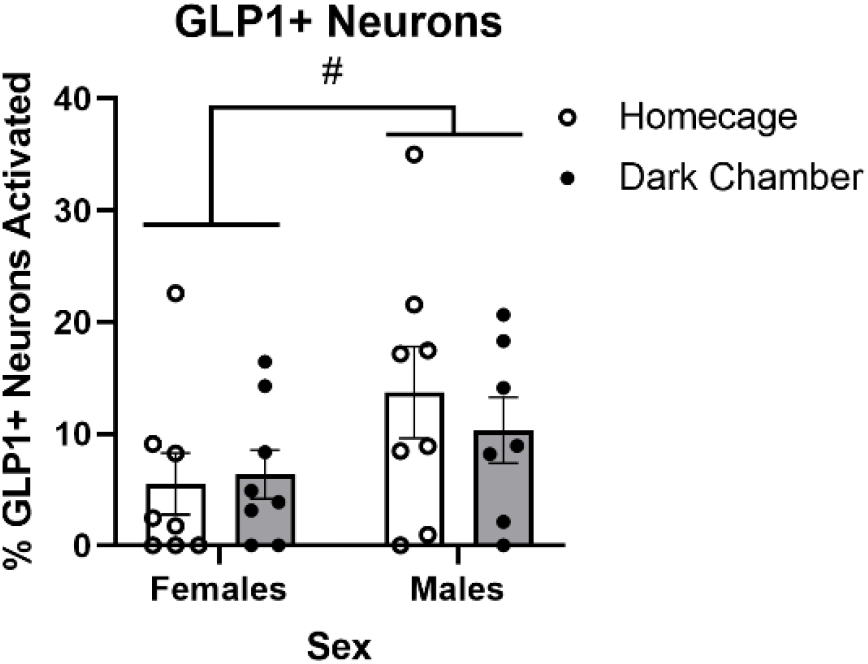
Re-exposure to the shock-paired dark chamber does not alter the percentage of GLP1+ neurons in the cNTS expressing cFos. #p < 0.1. Bars represent group mean±SEM.

**Figure 4.**
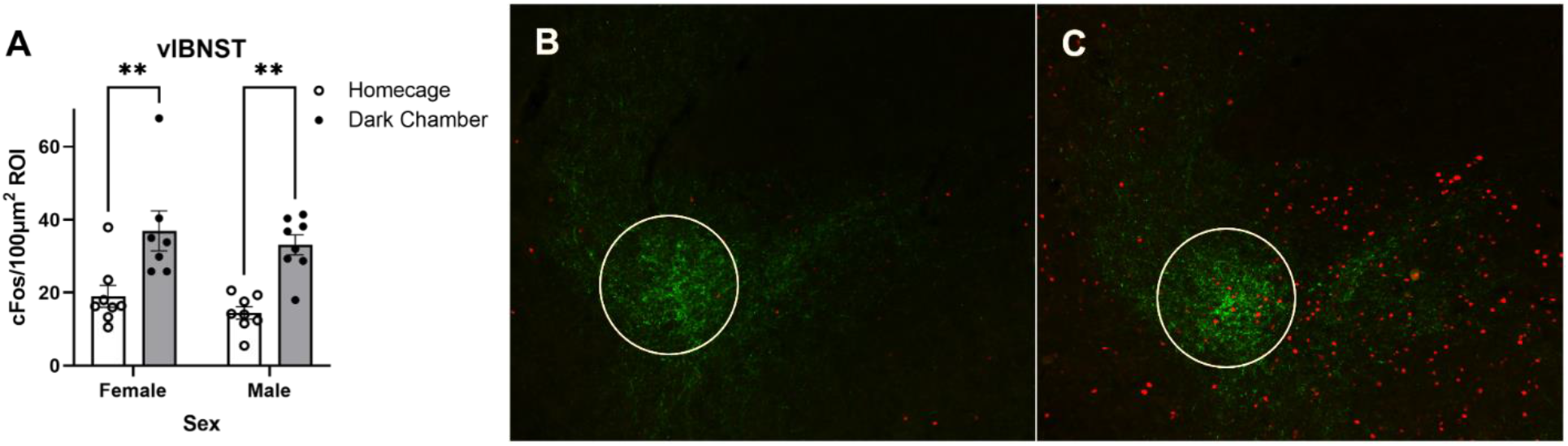
Re-exposure to the shock-paired dark chamber increases the average number of cFos+ neurons in the vlBNST. (A) Summary data reporting the average number of cFos+ neurons 100μm^2^ region of interest (ROI) in the vlBNST in dark chamber re-exposed rats vs. homecage controls (group mean±SEM). (B+C) Representative images depicting nuclear cFos (red) and DbH fiber and terminal (green) immunolabeling within the vlBNST from a homecage control rat (B) and a dark chamber re-exposed rat (C). A white circle outlines the ROI used for quantification of cFos. In A, **p < 0.01.

##### *PrRP*+ *Subpopulation*

Two-way ANOVA (Sex x Metabolic State) revealed a significant main effect of metabolic state on PrRP+ neuron activation in rats after dark chamber exposure [F(1,20) = 38.53; p < 0.0001], with no significant main effect of sex and no interaction between sex and metabolic state. Post-hoc Tukey’s multiple comparisons tests indicated a significant difference in PrRP neural activation between food deprived and *ad libitum* fed males (p = 0.0003) and females (0.0076) after dark chamber re-exposure.

#### 3.3.2. BNST

Two-way ANOVA (Sex x Metabolic State) revealed a significant main effect of metabolic state on the average number of cFos+ neurons within a 100μm^2^ ROI corresponding to the fusiform subnucleus of the vlBNST [F(1,21) = 9.304; p = 0.0061] in rats after dark chamber re-exposure, with no significant main effect of sex, and no interaction between sex and metabolic state. Post-hoc Tukey’s multiple comparisons tests indicated that food deprivation reduced the number of cFos+ neurons in the vlBNST in response to dark chamber re-exposure in both males (p = 0.0287) and females (p = 0.0287).

## 4. Discussion

### 4.1. Passive Avoidance Behavior

The present study demonstrates a contribution of metabolic state to the expression of conditioned avoidance behavior in male, but not female, rats. Specifically, we show that male rats tested under conditions of food deprivation more quickly entered the shock-paired dark chamber compared to rats tested under fed conditions, evidence for reduced expression of passive avoidance behavior during conditions of caloric deficit (Fig. 1). Conversely, metabolic state at the time of footshock training had no significant impact on the subsequent expression of conditioned passive avoidance. Previous research has demonstrated that being in a state of negative energy balance reduces innate anxiety-like behaviors in male rats (Maniscalco et al., 2015; Martin-Iverson & Stevenson, 2005). Here we extend these findings by demonstrating that negative energy balance also reduces avoidance behavior conditioned by aversive experience, and that this effect is sex-specific.

Examining the separate impact of food deprivation during training and testing phases allowed us to differentiate between potential effects of metabolic status on initial encoding/acquisition of fear memory vs. effects on later retrieval/expression of the memory. The present results indicate that metabolic state influences the expression but not the encoding of the fear memory underlying passive avoidance. In addition, passive avoidance latencies in rats that were trained and tested under matching metabolic state conditions (i.e., Ad Lib Tested/Ad Lib Trained & Food Dep Tested/Food Dep Trained) did not differ from avoidance latencies in rats with mismatched training and testing statuses (i.e., Ad Lib Tested/Food Dep Trained & Food Dep Tested/Ad Lib Trained), evidence that reductions in passive avoidance displayed by male rats tested under conditions of negative energy balance cannot be explained simply by viewing metabolic status as part of the testing and training contexts.

### 4.2 Neural Activation in Response to Dark Chamber Re-exposure

Quantitative analysis of cFos expression in rats exposed to the footshock-paired dark chamber revealed a potential contribution of A2 noradrenergic cNTS neurons in the expression of passive avoidance behavior, such that neural activation reflected sex- and metabolic state-specific avoidance behavior. Previous research has established that A2 neurons are activated in response to a variety of innate stressors that produce unconditioned avoidance responses (Maniscalco & Rinaman, 2018; Maruyama et al., 2001; Morales & Sawchenko, 2003; Rinaman, 2011). Here we demonstrate that A2 neurons, including the PrRP+ subpopulation, are activated in rats when they are re-exposed to the previously shock-paired dark chamber (Fig 2). Since the passive avoidance task used in our study relies on contextual stimuli present during training and testing, the task likely depends on neural circuits similar to those that underlie contextual fear conditioning (Izquierdo et al., 2016; Vazdarjanova & McGaugh, 1998; Whitlock et al., 2006). In this regard, our findings are consistent with previous research demonstrating that PrRP+ neurons in the cNTS are activated in rats exposed to contextual fear stimuli that were repeatedly paired with footshock (Yoshida et al., 2014; Zhu & Onaka, 2003). Our data extend those results by demonstrating that exposure only to a single, relatively mild footshock is sufficient to condition both passive avoidance behavior and neural activation.

Previous reports indicate that the ability of unconditioned acute stressors to activate A2 (including PrRP+) neurons in male rats is significantly attenuated by overnight food deprivation (Maniscalco & Rinaman, 2013; Maniscalco et al., 2015; Maniscalo et al., 2020). Here we newly demonstrate that overnight food deprivation reduces the ability of a conditioned contextual stressor (i.e., re-exposure to the shock-paired dark chamber) to activate A2 neurons, especially the PrRP+ subpopulation (Fig. 5B). The overall effect of food deprivation to reduce DbH+ neuron activation across 3 rostrocaudal levels of the cNTS (Fig. 5A) was driven by the largest effect for food deprivation to “silence” A2 neural activation at levels caudal to the AP (Fig. 5C left), the level at which the largest proportion of DbH+ A2 neurons express PrRP (Chen et al., 1999). While these findings do not provide causal evidence that PrRP+ neurons are involved in the expression of passive avoidance behavior, it is interesting to note the close parallel between effects of food deprivation on PrRP+ neuron activation and on passive avoidance behavior. Future studies will be necessary to determine whether PrRP neuronal activation is necessary and/or sufficient for full expression of this learned behavior. The mechanism by which food deprivation reduces A2/PrRP neuronal activation remains unclear, but potential mechanisms include deprivation-induced changes in vagal sensory feedback signaling from the gut (Appleyard et al., 2007). For example, circulating ghrelin levels increase during states of negative energy balance, ghrelin reduces the ability of vagal afferent stimulation to activate A2 neurons (Cui et al., 2011), and pharmacological antagonism of ghrelin receptor signaling partially rescues A2/PrRP neural activation in food-deprived rats (Maniscalco et al., 2020).

**Figure 5.**
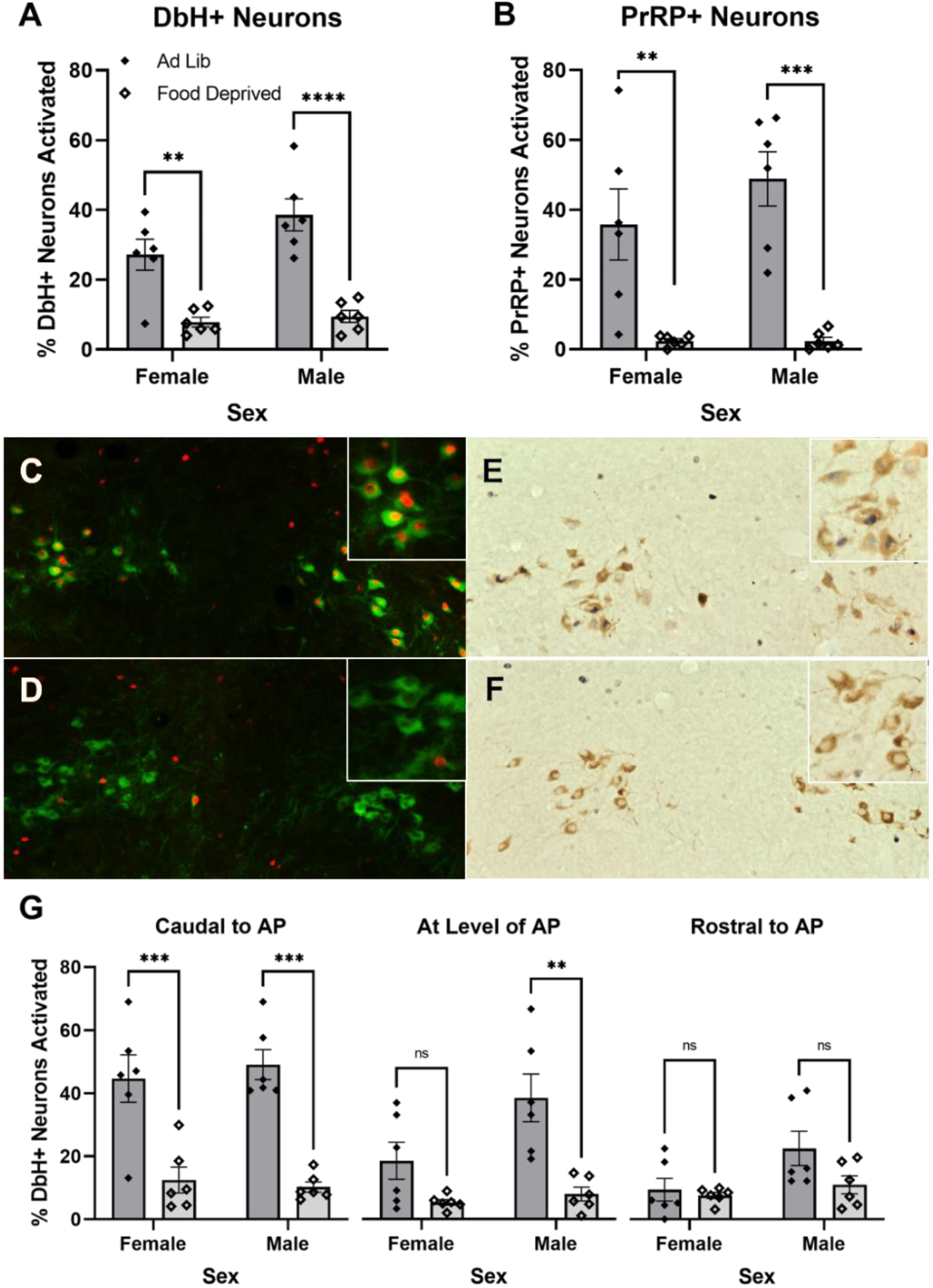
Overnight food deprivation reduces the percentage of noradrenergic cNTS neurons expressing cFos in rats re-exposed to the shock-paired dark chamber. (A) Food deprivation reduces activation of DbH+ neurons in both sexes. (B) Food deprivation reduces activation of the PrRP+ subset of noradrenergic neurons in both sexes. (C+D) Representative images depicting nuclear cFos (red) and cytoplasmic DbH (green) immunolabeling in a rat re-exposed to the dark chamber when *ad lib* fed (C), and in a rat that was food deprived during re-exposure (D). (E+F) Representative images depicting nuclear cFos (dark blue/black) and cytoplasmic PrRP (brown) immunolabeling in a rat re-exposed to the dark chamber when *ad lib* fed (E), and in a rat that was re-exposed while food deprived (F). Insets in the upper right corner of each panel (C-F) show immunolabeling at higher magnification. (G) Food deprivation reduces activation of DbH+ noradrenergic neurons in male and female rats. In females, this effect is specific to the most caudal level of the cNTS *(left)*. Food deprivation does not affect activation of DbH+ noradrenergic neurons rostral to the AP in either sex. In A, B, and G, ns=p > 0.05; **p < 0.01; ***p <0.001; ****p < 0.0001. Data in bar graphs represent group mean±SEM.

**Figure 6.**
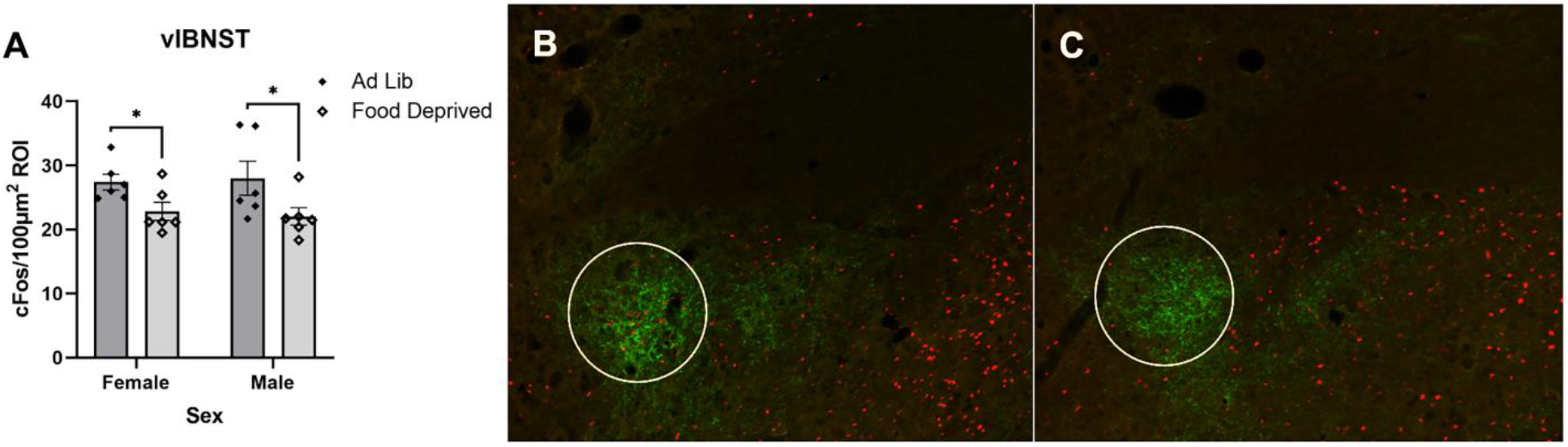
Overnight food deprivation reduces the average number of cFos+ neurons in the vlBNST in rats that were re-exposed to the shock-paired dark chamber. (A) Summary data reporting the average number of cFos+ neurons within a 100μm^2^ ROI in the vlBNST in female and male rats that were either *ad lib* fed or food deprived before chamber re-exposure. (B+C) Representative images depicting nuclear cFos (red) and DbH fiber and terminal immunolabeling (green) within the vlBNST from a rat that was *ad lib* fed (B) and a rat that was food deprived (C) prior to dark chamber re-exposure. A white circle outlines the ROI in which cFos was quantified. In A, *p < 0.05, and bars represent group mean±SEM.

A2/PrRP neurons densely innervate the fusiform subnucleus of the anterior vlBNST (Bienkowski & Rinaman, 2013; Maniscalco et al., 2015; Ricardo & Tongju Koh, 1978; Riche et al., 1990), and this limbic forebrain region displayed significantly more cFos activation in rats that were re-exposed to the dark chamber compared to activation in homecage controls (Fig. 4). Further, cFos activation within the vlBNST was reduced in rats that were food deprived before dark chamber re-exposure compared to activation in rats that were fed *ad libitum* (Fig. 6). Interestingly, this reduction in activation was specific to the ROI corresponding to the fusiform subnucleus and not the surrounding nuclei of BNST. These results are consistent with the idea that activation of A2/PrRP inputs to the vlBNST participates in the expression of passive avoidance. Noradrenergic signaling in the vlBNST has been broadly implicated in avoidance and other anxiety-like behavioral responses to innate stressors (Cecchi et al., 2002; Schmidt et al., 2019; Zheng & Rinaman, 2013), and BNST circuits are implicated in the expression of conditioned fear responses when rats are re-exposed to relevant contextual stimuli (Poulos et al., 2010; Zimmerman & Maren, 2011). A particularly interesting example supporting a role of noradrenergic signaling within the vlBNST in contextual conditioning is opiate withdrawal, which produces a robust conditioned place avoidance in rats (Aston-Jones et al., 1999). Opiate withdrawal activates A2 neurons and neurons within the vlBNST, and experimental reduction of noradrenergic signaling in the vlBNST reduces withdrawal-induced conditioned place avoidance (Aston-Jones et al., 1999). Results from the present study complement those findings, providing additional support for the idea that conditioned passive avoidance behavior depends on noradrenergic signaling in the vlBNST.

We were surprised to find that GLP1-expressing cNTS neurons were not activated by re-exposure to the dark chamber (Fig. 3). Central GLP1 signaling has been widely implicated in neuroendocrine and behavioral responses to acute innate stressors, including increased anxiety-like behavior (Anderberg et al., 2016; Kinzig et al., 2003; Huiyuan Zheng et al., 2019), and GLP1+ neurons are activated to express cFos in response to a variety of innate interoceptive and psychogenic stress stimuli (Holt & Rinaman, submitted; Maniscalco et al., 2015; Rinaman, 1999). Our new findings indicate that GLP1 neurons are not activated by conditioned stimuli associated with a prior stressor, whereas A2/PrRP neurons are activated both by innate and conditioned stressors. Interestingly, our data suggest that GLP1 neurons were even less activated by the conditioned context related to prior footshock experience in female rats compared to males (Fig. 3). It will be important to replicate and extend this observation in order to examine potential sex differences in GLP1 neural responses to other conditioned stress stimuli.

### 4.3 Sex Differences in Passive Avoidance Behavior

We found that male rats displayed more passive avoidance compared to similarly trained and tested female rats, consistent with previous literature (Beatty et al., 1973; Denti & Epstein, 1972; Drago et al., 1980; Van Oyen et al., 1979). This sex difference seems unrelated to differences in footshock sensitivity during training, given that female rats have lower shock response thresholds than males (Beatty & Beatty, 1970; Beatty & Fessler, 1977; Pare, 1969). It also is unlikely that the sex difference is related to perceived aversiveness of the light chamber, since male and female rats displayed similar latencies to enter the dark chamber during training before footshock (Fig. 1B), and previous studies report no sex differences in the movement of rats between light and dark chambers in non-punished (unshocked) tasks (Denti & Epstein, 1972; Võikar et al., 2001). Instead, the observed sex difference may be due to different behavioral strategies used by male and female rats during threat responses. Compared to males, female rodents tend to show less passive and more active responses to threatening stimuli (Archer, 1975; van Haaren et al., 1990), including less contextual freezing/immobility (Maren et al., 1994; Pryce et al., 1999), more escape behavior (Davis et al., 1976), and more active avoidance (Beatty & Beatty, 1970; Denti & Epstein, 1972). With this in mind, conditions of metabolic deficit may shift behavioral threat responses towards more active strategies, such that passive avoidance behavior is reduced in male rats to a greater degree than in females, which already display lower levels of passive avoidance. This idea would be consistent with early research demonstrating that food-deprived male rats display faster active escape from footshock (Amsel, 1950) and faster active avoidance latencies (Ley, 1965) compared to fed male rats.

Interestingly, similar sex differences exist in conditioned taste avoidance (CTA), another single-trial avoidance task in which rats avoid consuming foods or liquids that were previously paired with malaise-inducing or psychoactive drugs (Chambers et al., 1981; Chambers & Sengstake, 1976; King et al., 2015; Sherrill et al., 2011). While there is some inconsistency in the literature regarding sex differences in CTA, only studies in which rats were water deprived prior to testing (and thus indirectly food deprived, since water deprivation reduces food intake) failed to reveal sex differences in the expression of CTA (Roma et al., 2008; Vetter-O’Hagen et al., 2009). Indeed, results from studies directly examining the impact of water deprivation and sex on lithium chloride-induced CTA revealed that female rats display less CTA than male rats, unless rats are water deprived prior to testing (Chambers et al., 1981; Chambers & Sengstake, 1976). Water deprivation prior to testing does not impact CTA expression in female rats, whereas it reduces CTA in males to levels similar to females. Those results are consistent with our current finding that food deprivation reduces conditioned passive avoidance in male rats to a level roughly similar to that displayed by females.

### 4.4 Sex Differences in Neural Activation in Response to Dark Chamber Re-exposure

Activation of DbH+ neurons in rats after re-exposure to the shock-paired dark chamber differed by sex in a manner that depended on rostrocaudal level of the cNTS (Fig. 2C). Specifically, no sex difference was observed at levels rostral to the AP, whereas A2 neural activation at more caudal cNTS levels was more pronounced in male rats. These caudal cNTS levels correspond to the highest density of PrRP+ A2 neurons (Chen et al., 1999), which were activated to a greater extent in male vs. female rats after dark chamber re-exposure (Fig. 2B). While these observations do not demonstrate causality, it is interesting to note that females also displayed less passive avoidance behavior compared to males (Fig. 1).

We did not evaluate potential effects of female estrous cycle on either behavior or neural activation, but this could be explored in future investigations. Compared to males, female rats display higher levels of estrogen receptors within the cNTS (Schlenker & Hansen, 2006), expression of which is influenced by both stress and food deprivation (Estacio, Tsukamura, et al., 1996; Estacio, Yamada, et al., 1996; Reyes et al., 2006). Some studies report that PrRP mRNA expression within the cNTS varies across the estrous cycle (Anderson et al., 2003; Feng et al., 2007; Kataoka et al., 2001). Additionally, estrogen signaling increases vagal afferent signaling to the cNTS (Ciriello & Caverson, 2016) and has been reported to modulate activation of cNTS neurons in response to other stimuli (Asarian & Geary, 2007; De Oliveira et al., 2003).

### 4.5 Conclusions

In summary, results from the present study demonstrate that a moderate state of negative energy balance reduces conditioned passive avoidance behavior in male, but not female, rats. We further demonstrate that re-exposure to a conditioned aversive context activates PrRP+ A2 noradrenergic neurons and their downstream targets within the vlBNST, and that this neural activation is reduced by food deprivation. These novel findings support the view that A2 noradrenergic signaling to the vlBNST contributes to the expression of conditioned passive avoidance behavior. This ascending neural signaling pathway provides a potential mechanism by which conditioned avoidance behavior is influenced by interoceptive sensory feedback from the body and may suggest new strategies for the treatment of excessive avoidance in anxiety and stress-related disorders.

